# The Protein Phosphatase 4 complex promotes the Notch pathway and *wingless* transcription

**DOI:** 10.1101/113852

**Authors:** Eric T. Hall, Tirthadipa Pradhan-Sundd, Faaria Samnani, Esther M. Verheyen

## Abstract

The Wnt/Wingless (Wg) pathway controls cell fate specification, tissue differentiation and organ development across organisms. Using an *in vivo* RNAi screen to identify novel kinase and phosphatase regulators of the Wg pathway, we identified subunits of the serine threonine phosphatase Protein phosphatase 4 (PP4). Knockdown of the catalytic and the regulatory subunits of PP4 cause reductions in the Wg pathway targets Senseless and Distal-less. We find that PP4 regulates the Wg pathway by controlling Notch-driven *wg* transcription. Genetic interaction experiments identified that PP4 likely promotes Notch signaling within the nucleus of the Notch-receiving cell. Although the PP4 complex is implicated in various cellular processes, its role in the regulation of Wg and Notch pathways was previously uncharacterized. Our study identifies a novel role of PP4 in regulating Notch pathway, resulting in aberrations in Notch-mediated transcriptional regulation of the Wingless ligand. Furthermore, we show that PP4 regulates proliferation independent of its interaction with Notch.

**Summary statement:** The protein phosphatase 4 complex promotes Notch signaling and target gene expression during Drosophila wing development.

## Introduction

The progression from a fertilized egg into a multicellular organism is a complex process, requiring proliferation and intricate cell-cell communication between individual cells for the eventual formation of tissues and organs. Only a handful of evolutionarily conserved signal transduction pathways are used reiteratively, both spatially and temporally to control development. In metazoans, the Wnt signaling [Wingless (Wg) in Drosophila] pathway regulates growth and proliferation, cell-fate differentiation, stem-cell renewal and homeostasis (Clevers and Nusse, 2012; Swarup and Verheyen, 2012). Wnt signaling alone does not control all these processes; its activity is extensively regulated by other signaling pathways and cellular mechanisms (Collu et al., 2014; Itasaki and Hoppler, 2010; Kim and Jho, 2014; Zeller et al., 2013). Determining how these interactions occur is critical for understanding basic cellular function and disease progression, as the disruption of the Wnt pathway has been implicated in a variety of developmental disorders and cancer (Clevers and Nusse, 2012).

The Drosophila wing imaginal disc is a powerful tool for studying Wg signaling (Swarup and Verheyen, 2012). In the developing wing disc the Wg ligand is expressed throughout different stages of disc development. At the end of the larval third instar stage, Wg expression is confined to the presumptive wing margin along the dorsal/ventral (D/V) boundary, which controls patterning and fate specification (Couso et al., 1994; Williams et al., 1993). Wg produced in this narrow band of cells is secreted and acts as a morphogen to induce the nested expression of target genes including Senseless (Sens) and Distal-less (Dll), in the flanking non-boundary cells (Neumann and Cohen, 1997; Zecca et al., 1996).

The directed expression of Wg at the D/V boundary requires the transmembrane receptor Notch in these boundary cells. The Notch ligands Delta (Dl) and Serrate (Ser) signal from the flanking non-boundary cells, inducing proteolytic cleavages of Notch to generate a free Notch intracellular domain (N^ICD^) (Bray, 2016; Fortini, 2009). N^ICD^ translocates to the nucleus where it binds transcriptional co-activators and DNA binding proteins to initiate target gene transcription, including *wg* and *cut* (de Celis et al., 1996; Rulifson and Blair, 1995). The absence of Notch results in reduced *wg* transcription and therefore reduced Wnt pathway activation (Rulifson and Blair, 1995).

Both Notch and Wg signaling act to regulate common developmental processes such as tissue patterning, fate specification and growth of different Drosophila appendages (Hing et al., 1994). These two pathways share a number of common regulators which affect the activity of their signaling outcome. In an *in vivo* RNAi screen to identify novel kinase and phosphatase modulators of the Wg pathway, we found that the components of the Protein Phosphatase 4 (PP4) complex appeared to promote Wg signaling (Swarup et al., 2015). The serine threonine phosphatase PP4 belongs to the Protein Phosphatase 2A (PP2A) group of phosphatases (Cohen et al., 2005). Similar to what is found with PP2A, PP4 forms a heterotrimeric complex, which in Drosophila consists of a catalytic subunit, Protein Phosphatase 4-19C (PP4-19C), and two regulatory subunits called Protein Phosphatase 4 Regulatory subunit 2-related protein (PPP4R2r) and PP4R3/Falafel (Flfl) (Cohen et al., 2005; Gingras et al., 2005).

PP4 is a highly conserved phosphatase seen across metazoans, and has been implicated in a wide range of cellular processes, including chemotaxis in slime molds (Mendoza et al., 2007), developmental signaling pathways such as Hedgehog (Jia et al., 2009), JNK (Huang and Xue, 2015; Zhou et al., 2002), Insulin-like growth factor (Mihindukulasuriya et al., 2004), as well as TOR (Raught et al., 2001). The major functional role of PP4 is as a key regulator in cell cycle progression and regulation of cell division (Helps et al., 1998; Huang et al., 2016). No previous studies have implicated PP4 in Notch or Wnt/Wg signaling.

In this study we demonstrate that our previous observations of reduced Wg signaling due to knockdown of PP4 components are due to effects on *wg* transcriptional regulation by Notch. Using genetic interaction studies and expression of mutant Flfl, we determine that PP4 promotes the activity of nuclear Notch. We further elucidated that the function of PP4 in promoting Notch signaling was independent of its previously described role in cell cycle progression and proliferation. Taken together we have identified a novel role for PP4 in promoting Notch signaling and expression of *wg* during Drosophila development.

## Results

### PP4 promotes Wg signaling in the Drosophila wing imaginal disc

In a screen for modifiers of Wg signaling in the Drosophila wing imaginal disc, three components of PP4 were found to reduce Wg target genes following their knockdown through RNAi (Swarup et al., 2015). An involvement of PP4 in Wg signaling has not been previously identified, so we were curious to determine mechanistically how PP4 may be involved in regulating the output of the Wg pathway. In the developing wing imaginal disc the Wg target genes *Sens* and *Dll* are expressed in a distinct nested pattern along the dorsoventral (D/V) boundary (Fig. 1A, A’). We utilized *dpp-GAL4* expressed in a narrow band of cells along the anterior/posterior (A/P) boundary (marked by GFP; Fig. 1B’, B”) to express RNAi constructs to knockdown expression of the individual PP4 components. The knockdown of the catalytic subunit *PP4-19C* caused a strong reduction in Sens and Dll (Fig. 1C, C’). Reduction of the PP4 targeting subunits *ppp4R2* or *flfl* via RNAi caused reduced Dll protein levels (Fig. 1E’, G’), but did not noticeably affect Sens (Fig. 1E, G).

**Figure 1.**
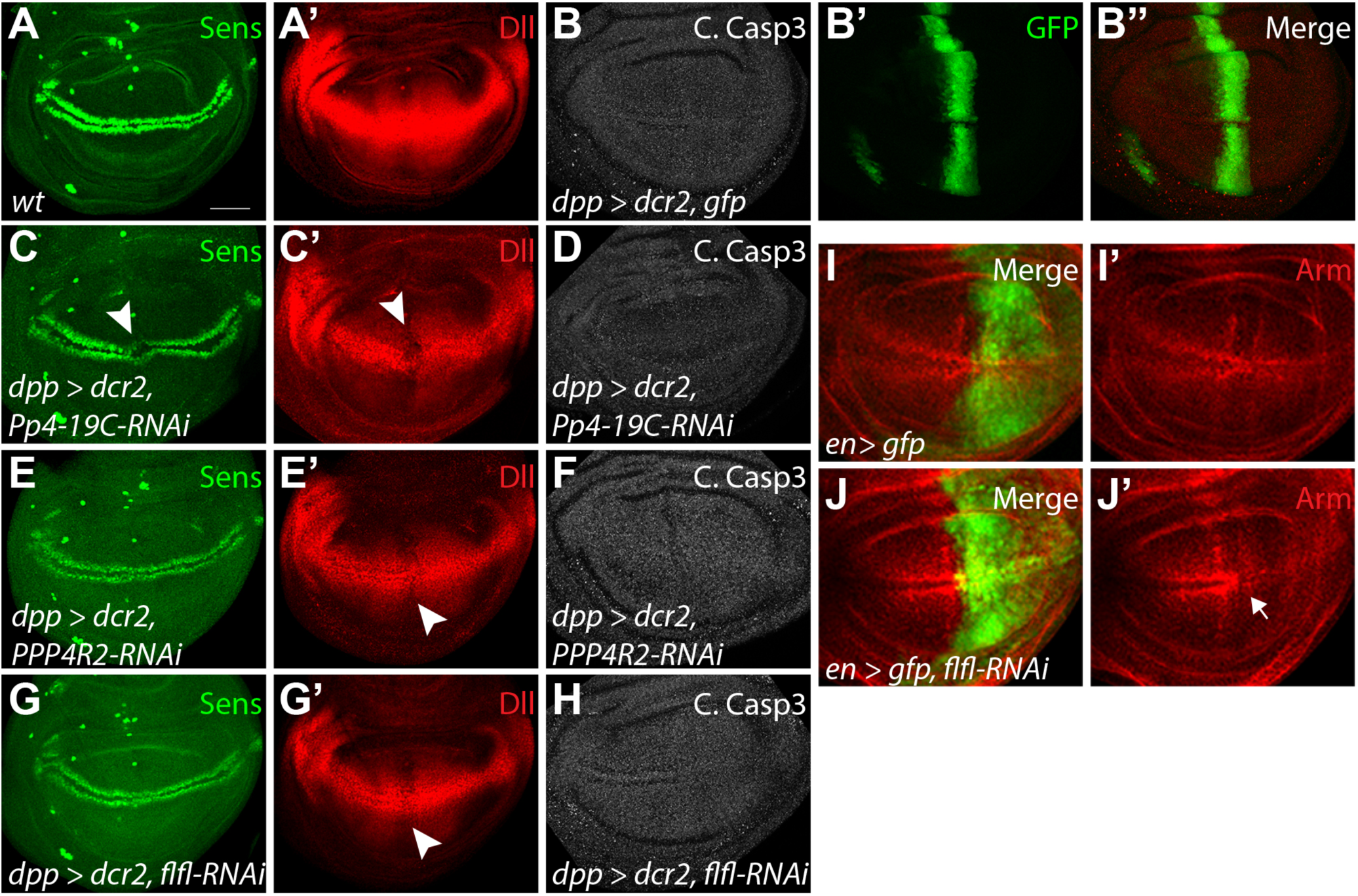
Reduction of PP4 subunits inhibits Wg pathway activation without inducing cell death. (A-B’’) Wild-type expression pattern of Wg target genes Sens (A), and Dll (A’), as well as cleaved caspase 3 (B) with normal expression pattern of *dpp-Gal4* (B’, B”) along the anterior/posterior boundary in the developing wing disc. (C-D) The knockdown of PP4-19C with RNAi along the anterior/posterior boundary causes a loss of Sens (C, arrow head) and Dll (C’, arrow head), but does not elevate C. Caspase 3 activity (D). (E-F) Knockdown of PPP4R2 was able to effectively reduce Dll (E’, arrow head), but did not affect Sens (E) or C. Casp3 (F). (G-H) Flfl knockdown did not affect Sens (G) or C. Casp3 levels (H), but did reduces Dll (G’, arrow head). Normal expression pattern of *en-Gal4* (I) in the posterior domain of the wing disc, shown with Arm (I, I’). (J-J’) The knockdown of Flfl in the posterior domain (shown with GFP) induced a marked reduction in stabilized Arm (J’). Scale bar: 50 μm.

To confirm that the reduction of PP4 components did not critically affect cell viability as loss of PP4 has been shown to promote JNK-dependant cell death (Huang and Xue, 2015), discs were stained for the apoptotic marker, cleaved caspase-3 (C. Casp-3). Compared to control cells expressing GFP (Fig. 1B, B”), reduction of any individual PP4 component did not increase levels of apoptosis within the *dpp* domain of the imaginal disc (Fig. 1D, F, H). Following Wg pathway activation, the key effector protein Arm is stabilized at the highest concentration in two bands of cells flanking the Wg-producing cells of the D/V boundary (Fig. 1I, I’) (Peifer et al., 1991). Expression of *flfl-RNAi* in the posterior domain of the wing imaginal disc, using *engrailed (en)- Gal4* (Fig. 1I, J), caused a reduction of stabilized Arm (Fig. 1J, arrow in J’). In subsequent experiments, we utilized *flfl-RNAi* to reduce PP4 activity, as it has been previously confirmed as a functional indicator of the entire complex (Sousa-Nunes et al., 2009). Together, these data suggest PP4 is required for promoting Wg pathway activation.

### PP4 promotes Wg signaling through Notch pathway activation

As the reduction of Wg target genes and Arm was apparent upon knockdown of PP4 components, we next wanted to look at the Wg ligand and its transcription. In third instar wing imaginal discs, *wg* is transcribed, translated, and undergoes post-translational modification, which can affect its stability, before being secreted to activate the Wg pathway in neighbouring cells (Franch-Marro et al., 2008; Tanaka et al., 2002). Utilizing Wg antibodies, and the *wg* transcriptional reporter, *wg-lacZ,* we could identify any defects in the ligand’s transcription, processing, or stability. In a wild type wing imaginal disc, Wg and *wg-lacZ* expression are refined along the D/V boundary in a narrow band 2-3 cells wide (Fig. 2A, B). Expression of *flfl-RNAi* in the posterior domain of the wing disc, using *hedgehog (hh)-Gal4,* resulted in a reduction of both total Wg protein levels and transcription (Fig. 2E, F) suggesting PP4 is involved in regulation of *wg* transcription.

**Figure 2.**
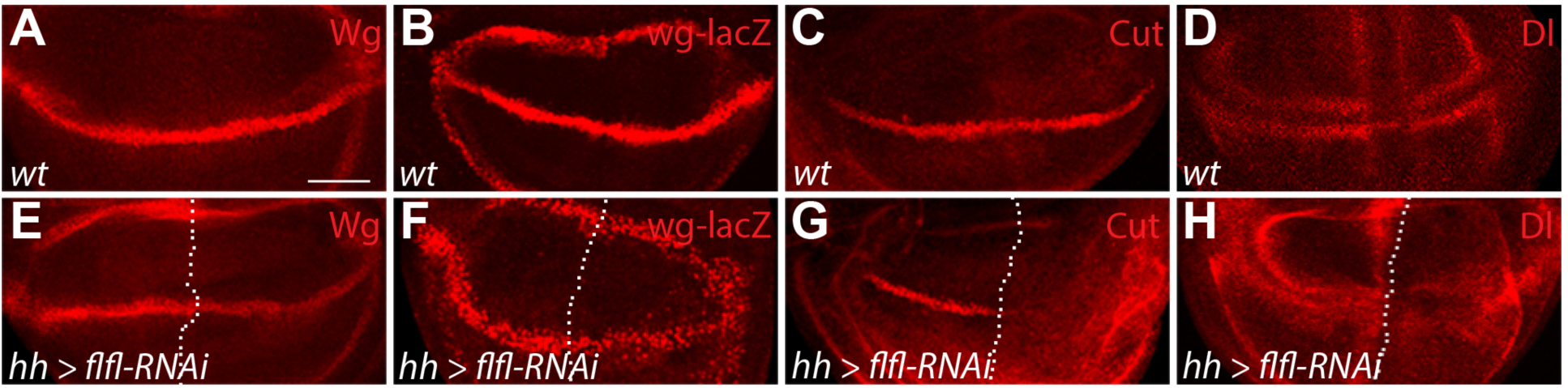
PP4 promotes Wg signaling through Notch pathway activation. (A-D) Wild-type pattern of Wg protein (A), *wg* transcription (B), Cut protein (C), and Dl protein (D) in the developing wing disc. (E-H) Using *hh-Gal4,* expressed in the posterior domain of the wing disc (right of the dotted line), to express *flfl-RNAi,* caused a reduction in total Wg protein levels (E), as well as *wg* transcription (F), and induced a loss of Cut (G), and Dl (H). Scale bar: 50 μm.

*wg* transcription is controlled along the D/V boundary of the wing disc by Notch signaling (Rulifson and Blair, 1995). We next wished to determine if PP4 regulation of *wg* transcription is mediated through the involvement of the Notch signaling pathway. *cut*, another Notch target gene (de Celis et al., 1996), is expressed in a similar pattern as Wg along the D/V boundary (Fig. 2C). The reduction of *flfl* in the posterior domain of the wing disc via RNAi resulted in a strong loss of Cut expression, indicating an overall reduction in Notch signaling (Fig. 2G). Looking at the Notch ligand Delta (Dl) which is enriched in the cells adjacent to the D/V boundary (Fig. 2D), it was apparent that *flfl-RNAi* expressed in the posterior domain of the disc, resulted in reduced Dl and a failure of its refinement (Fig. 2H). We could not discern if this effect on Dl is from upstream regulation of *Dl* expression, or on Notch activation itself, as the refinement of Dl involves a cis/trans feedback mechanism with N for pattern refinement, through lateral inhibition of each gene (Axelrod, 2010). Together, these results demonstrate that PP4 normally appears to influence Notch signaling to promote multiple pathway targets including *wg*.

### PP4 promotes Notch signaling in the Notch signal receiving cells

Having identified that Flfl, and by extension PP4, is involved in promoting Notch signaling we sought to further elucidate how. During wing imaginal disc development Notch and its ligands Dl and Serrate (Ser) undergo refinement through lateral inhibition resulting in high levels of active Notch (N^ICD^) being expressed along the D/V boundary and suppressed in the flanking cells (Fig. 3D arrow head), which conversely have high levels of Dl (Fig. 2D) and Ser (de Celis et al., 1996). This feed forward loop of lateral inhibition creates Notch signal-sending cells (cells flanking the D/V boundary), and signal-receiving cells (D/V boundary cells) with active Notch signaling (Axelrod, 2010).

**Figure 3.**
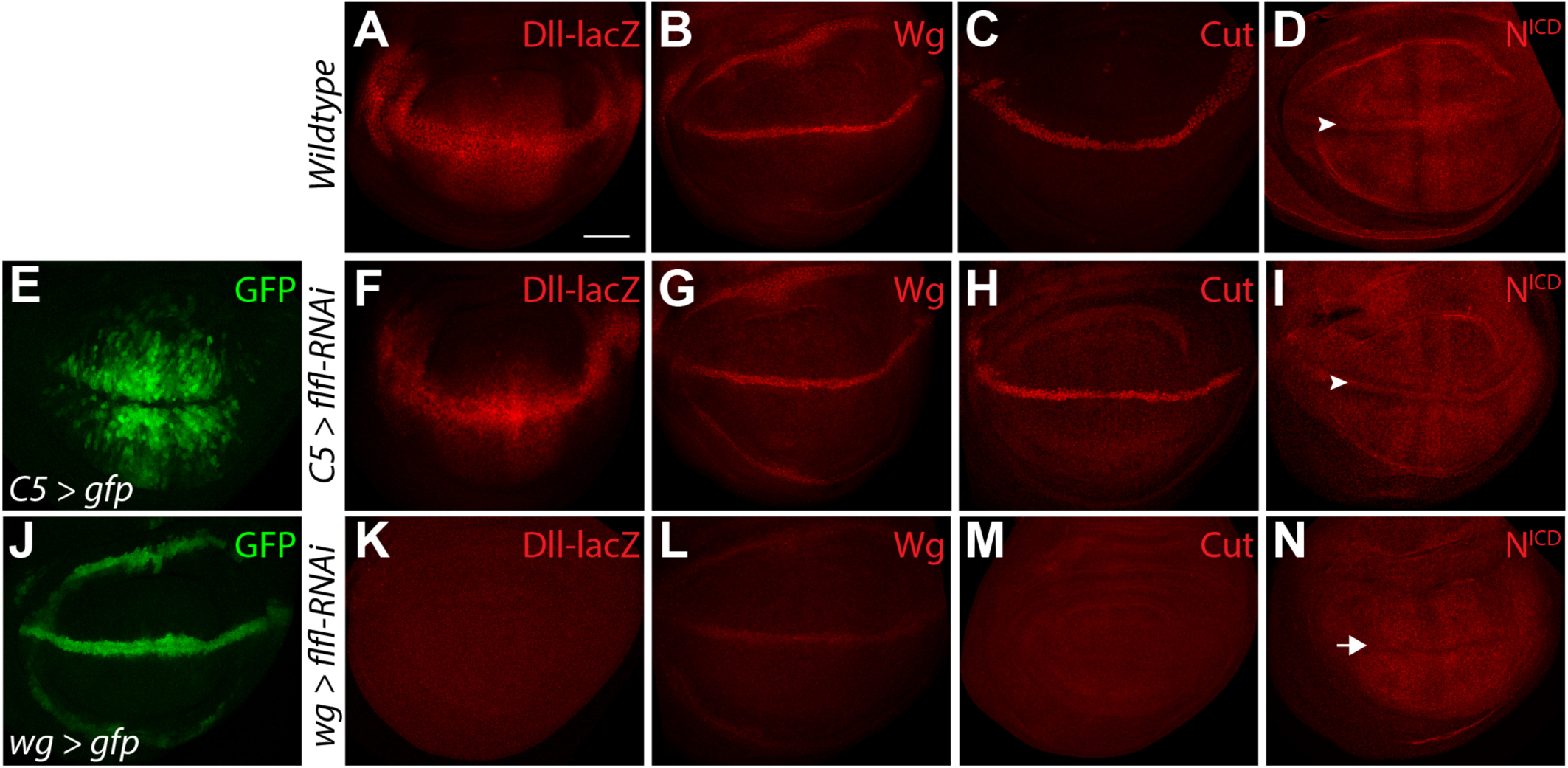
PP4 promotes Notch signaling in the Notch-signal receiving cells. (A-D) Wild-type expression pattern of *Dll-lacZ* (A), Wg (B), Cut (C), and N^ICD^ (D) in the developing wing disc. N^ICD^ is enriched along the dorsal/ventral (D/V) boundary (D, arrowhead) and suppressed in the adjacent cells. (E) Expression pattern of *C5-Gal4* driving GFP in the D/V boundary flanking cells of the wing pouch. (F-I) The knockdown of *flfl* with RNAi in the D/V boundary flanking cells does not affect *Dll-lacZ* (F), Wg (G), Cut (H), or N^ICD^ (I). (J) Expression pattern of *wg-Gal4* driving GFP along the D/V boundary. (K-N) Knockdown of *flfl* in the D/V boundary cells causes a loss of *Dll-lacZ* (K), strong reduction of Wg (L), loss of Cut (M), and a failure of N enrichment along the D/V boundary (N, arrow). Scale bar: 50 μm.

To further analyze the role of the PP4 complex in the complementary ligand-expressing and Notch-expressing cells, we used *C5-Gal4* and *wg-Gal4* respectively, to express *flfl-RNAi* in the wing imaginal disc. *C5-Gal4* is expressed in N ligand expressing cells flanking the D/V boundary (Fig. 3E), while *wg-Gal4* is expressed along the D/V boundary, in the active N signal receiving cells (Fig. 3J). By looking at Wg and Notch target genes, we could determine in which cells Flfl and by extension PP4 is working to affect the Notch pathway. *C5>flfl-RNAi* appeared to have no affect on *Dll* expression (Fig. 3A, F), Wg (Fig. 3B, G), or Cut (Fig. 3C, H). The enrichment of the N^ICD^ along the D/V boundary also appeared to be unaffected (Fig. 3D, I arrowheads). *wg>flfl-RNAi* gave very contrasting results. Knockdown of *flfl* in the Notch receptor expressing cells resulted in a complete loss of *Dll-lacZ* (Fig. 3K), strong reduction of Wg (Fig. 3L), loss of Cut (Fig. 3M), and a failure of enrichment of N^ICD^ (Fig. 3N). Knockdown of the other components of PP4 using *wg-Gal4*, but not *C5-gal4*, also caused loss of Wg expression (Fig. S1A-D). Taken together, these results suggest that PP4 functions within the Notch-expressing cell to promote full pathway activation and target gene expression.

### PP4 functions within the nucleus to promote Notch signaling

To further refine where PP4 functions within the Notch signal receiving cell, we utilized mutant transgenes of Flfl and N in the adult Drosophila wing. During pupal wing metamorphosis, the activation and refinement of Dl and N are required to refine adult vein formation (Huppert et al., 1997). Notch signaling is also critical for the development of sensory bristles along the adult wing margin (reviewed in Posakony, 1994). Any developmental defects from expression of the various Flfl transgenes in adult wings could provide insight into PP4’s role in the Notch pathway. Using *MS1096-Gal4,* which is expressed across the entire developing wing pouch, to ectopically express wild type Flfl or a cytoplasmic form, Flfl ^Δ 3NLS+2NES^ (Flfl-cyto), had no effect on the adult wing compared to wild type (Fig. 4A-C). Endogenous Flfl is a predominantly nuclear protein and the wild type Flfl transgene has been shown to function similarly (Sousa-Nunes et al., 2009). Knockdown of *flfl* via RNAi induced ectopic and thicker veins in the adult wing (Fig. 4D), a hallmark of reduced Notch activity (Huppert et al., 1997). This phenotype could be suppressed by reintroduction of the wild type *flfl* transgene (Fig. 4E), but not with expression of *flfl-cyto* (Fig. 4F). This suggests that Flfl functions within the nucleus, rather than the cytoplasm, to promote Notch.

**Figure 4.**
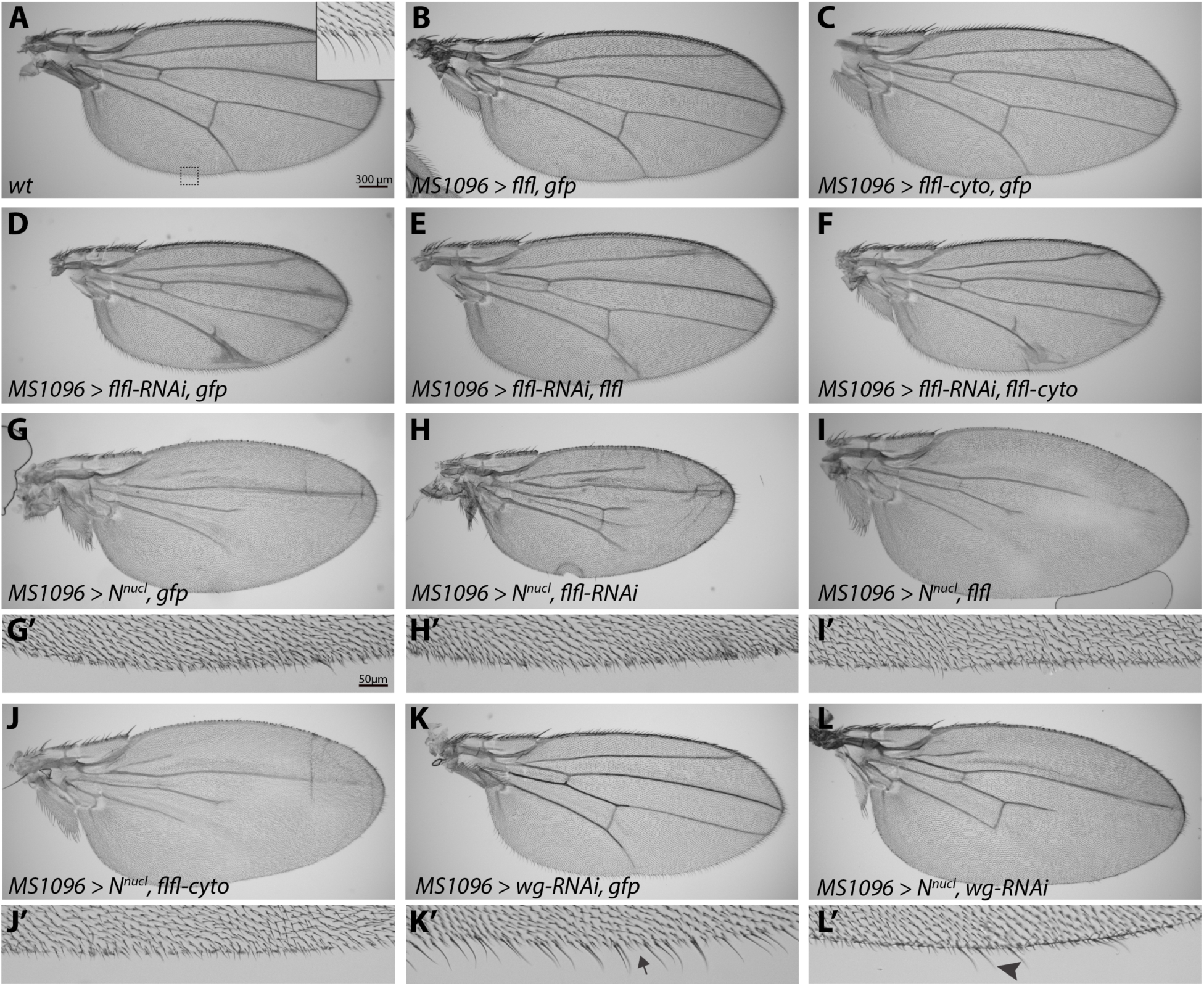
PP4 likely functions within the nucleus to promote Notch. (A-C) Adult wild-type wing and margin (A, inset). Over expression of Flfl (B), or Flfl-cyto(C) throughout the entire wing does not induce any noticeable phenotype. (D-E) Knockdown of Flfl induces ectopic veins and thickening of veins, as well as a reduced wing size (D). This effect can be primarily rescued by reintroduction of a full length Flfl transgene (E), but not by Flfl-cyto (F). (G-L’) Overexpression of N^nucl^ induces a loss of wing veins (G) and wing margin bristles (G’). Knockdown of *flfl* induced a mild rescue the N^nucl^ loss of vein phenotype, and still reduces the overall wing size (H, H’). The over expression of Flfl (I, I’), or Flfl-cyto (J, J’) did not disrupt the N^nucl^ phenotype. (K-L’) Expression of *wg-RNAi* in the wing induced sporadic loss of margin bristles (K, K’ arrow). N^nucl^ with *wg-RNAi* is able to maintain several margin bristles (L, L’ arrowhead).

To confirm the hypothesis that Flfl likely acts in nuclear Notch signaling, we expressed a construct encoding the intracellular domain of Notch that localizes to the nucleus (*N^nucl^*)(Rebay et al., 1993). This activated nuclear Notch suppressed wing vein formation and the formation of sensory bristles (Fig. 4G, G’) compared to wild type (Fig. 4A). Notch-dependent activation of *wg* expression is essential for Wg signaling to induce expression of proneural genes like *Sens* for the specification of sensory organ precursor (SOP) cells (Nolo et al., 2000). SOPs then divide and differentiate giving rise to the sensory bristles in the adult fly via Notch signaling (Guo et al., 1996; Hartenstein and Posakony, 1990). Although ectopic Notch signaling increases the number of SOPs in the wing disc via Wg, they do not differentiate correctly resulting in double socket cells, and loss of bristle cells (Guo et al., 1996). Knockdown of *flfl* in the *N^nucl^* expressing cells was able to partially recover vein loss, but did not significantly rescue the bristle defect (Fig. 4H arrow, H’). Conversely, the expression of Flfl or Flfl-cyto had no affect on the *N^nucl^* phenotype (Fig. 4I-J’).We interpret the inability to specifically rescue the bristle defect as being due to the combination of the strength of N^nucl^ as well as an incomplete knockdown of *flfl* via RNAi. To confirm that this phenotype could be rescued in our assay, we tested whether loss of *wg* could rescue the effect since up-regulated target genes expression causes the *N^nucl^* phenotype. Expression of a weak *wg-RNAi* transgene was able to induce sporadic sensory bristle loss (Fig. 4K, K’ arrow), due to reduced SOPs (Nolo et al., 2000; Parker et al., 2002). When combined: with *N^nucl^, wg-RNAi* can partially suppress the overactive Notch phenotype of inhibited bristle formation (Fig. 4L, L’ arrowhead).

### Flfl is required for proliferation and maintenance of overall tissue size independent of Notch signaling

Adult flies with reduced *flfl* expression displayed smaller wing blades compared to control flies (Fig. 4A, D). This was expected given PP4’s known role in cycle progression and growth (Helps et al., 1998; Huang et al., 2016; Martin-Granados et al., 2008; Zhuang et al., 2014). Notch has been implicated in cell proliferation in the wing imaginal disc, but a direct mechanism for its involvement is not fully understood (Baonza and Garcia-Bellido, 2000; Giraldez and Cohen, 2003; Go et al., 1998). We quantified the area of the adult wings of the different genotypes, in order to determine if PP4/Flfl’s role in growth is mediated through Notch signaling. Overexpression of Flfl and Flfl-cyto had no significant effect on wing size compared to wild type (Fig. 5A, box plot a,b,c). Knockdown of *flfl* resulted in a ~28% reduction in wing size, and could be fully rescued by the wild type *flfl* transgene (Fig. 5A, box plot d,e). Flfl-cyto; was able to slightly rescue the growth defect from *flfl-RNAi,* but not to a significant level (Fig. 5A, box plot f). The ability to rescue partially may be due to PP4’s role in mitotic progression after nuclear envelope break down at prometaphase, allowing for Flfl-cyto to perform its function at this step in mitosis (Huang et al., 2016). Wings expressing N^nucl^ did not have a significantly smaller area than wild type wings, and were unable to rescue the growth defect from *flfl-RNAi* (Fig. 5A, box plot g,h). As N^nucl^ is unable to rescue the growth defect from reduced Flfl levels, it indicates that PP4/Flfl’s role in regulating growth is likely independent to its function in propagating Notch signaling. An alternative interpretation could be that Flfl acts downstream of N^nucl^ or that Flfl is required for proper Notch function in the proliferation.

**Figure 5.**
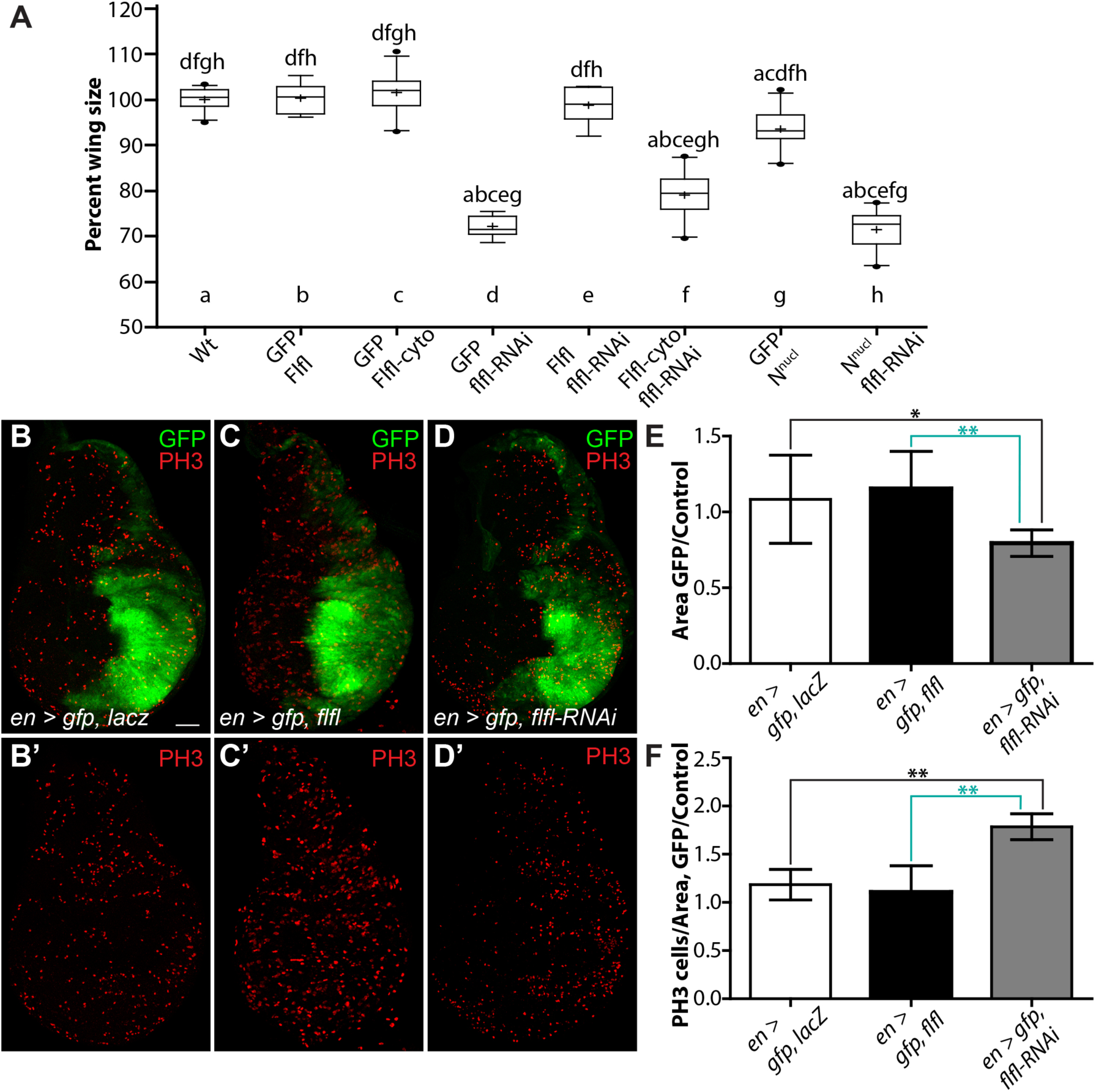
Flfl is required for proliferation and overall tissue size independent of Notch signaling. (A) Box plots representing total wing area of genotypes shown in Fig. 4 (n = 8-13). Over expression of Flfl (b) or Flfl-cyto (c) did not affect wing size compared to wild type (a). *flfl-RNAi* caused a significant reduction in wing size (d). The *flfl-RNAi* size defect could be fully rescued by reintroduction of full length Flfl (e), but no effect was seen with Flfl-cyto (f). Wings expressing N^nucl^ (g) are slightly smaller than wild type wings. N^nucl^ and *flfl-RNAi* wings (h) have a significant size reduction compared to wild type (a), equivalent to that of *flfl-RNAi* alone (d). Data are presented as box plot 25-75 percentile, whiskers 10-90 percentile, (—) median, (+) mean, (•) outliers, letters above representing significance from corresponding genotypes, (*p<0.01*) generated from one-way ANOVA. (B-F) The normal expression pattern of *en-Gal4* marked by GFP (B) in the posterior domain of the developing wing disc, shown with mitotic cell marker PH3 (B’) and represented as a ratio of posterior domain vs. the anterior control (n = 8)(E, F). Over expression of Flfl in the posterior domain had no effect on area (C, E) or proliferation rate (n = 7)(C’ F). The knockdown of Flfl with RNAi in the posterior domain induced a significant reduction in area (D, E), and exhibited a significantly higher number of PH3 positively marked cells (n=9)(D’ F). Data are presented as means ± s.d., (*)*p<0.05,* (**)*p<0.01* generated from one-way ANOVA. Scale bar: 50 μm.

To determine if the growth defects from loss of PP4 were due to decreased cell proliferation or overall cell size, we looked at the developing wing imaginal disc. We utilized *en*-*Gal4* driving GFP to mark the posterior portion of the wing disc, representing roughly 50% of the overall tissue (Fig. 5B). We compared the size of the GFP-positive region to the control anterior side of the disc in different genotypes to determine the effects that changes in the levels of Flfl have on tissue growth. In control discs expressing *UAS-lacZ,* the ratio between the internal control area of the anterior, to the GFP positive posterior was equal (Fig. 5B, E). Overexpression of Flfl in the posterior domain did not affect the posterior/anterior area ratio (Fig. 5C, E). However, reduction of *flfl* via RNAi resulted in a significant decrease in size of the posterior domain (Fig. 5D, E).

We simultaneously looked at the number of mitotic cells in these genotypes using a phospho-Histone H3 (Ser10) (PH3) antibody. There was no significant difference between our control discs and those overexpressing of Flfl (Fig. 5B’, C’, F). Surprisingly, we found that *flfl-RNAi* tissue had significantly elevated levels of PH3 positive cells (Fig. 5D’, F). This result was perplexing considering the decreased tissue size, yet apparent increase in proliferation rate. As PH3 marks condensed chromatin prior to chromosomal segregation in cells along the G2/M transition (Hendzel et al., 1997), it is possible that *flfl-RNAi* cells were arrested during early mitosis, and had not undergone mitotic exit. Previously, Sousa-Nunes et al. (2009)identified a similar effect. Drosophila *flfl/+* brains exhibited much lower rates of Bromodeoxyuridine (BrdU) incorporation and reduced proliferation, yet exhibited elevated levels of PH3 positive cells, demonstrating *flfl* is important for mitotic progression. Similar results have been found in multiple cases (Huang et al., 2016; Martin-Granados et al., 2008). Our results suggest a similar function for PP4/Flfl in the wing disc, where PP4 is critical for cell cycle progression and mitosis affecting proliferation rates and overall tissue size, yet this function is independent of its role in promoting Notch signaling.

### aPKC is not involved with PP4 and Notch signaling in the Drosophila wing imaginal disc

We examined whether PP4 mechanistically interacted with other known modulators of the Notch pathway. One well-characterized regulator is atypical protein kinase C (aPKC). aPKC is a protein kinase widely studied for its role in developmental processes, including asymmetric cell division (ACD). The regulators of ACD (including aPKC, Bazooka, and Crumbs) act upstream of Notch signaling and determine the identity of the Notch signal sending and signal receiving cells (reviewed in Knoblich, 2008). aPKC also promotes the Notch pathway by inhibiting Numb-mediated endocytosis of the Notch pathway components (Frise et al., 1996; Smith et al., 2007; Wang et al., 2006). Previous studies have shown a role for PP4/Flfl in the localization of the Miranda complex to promote neuroblast ACD in Drosophila by acting downstream or parallel to aPKC (Sousa-Nunes et al., 2009). Since such a role has not been identified in a proliferating epithelium like the wing disc we sought to investigate if this mechanism was conserved during the development of the wing disc.

Using *MS1096-Gal4* to express *aPKC-RNAi* in the wing disc resulted in small, crumpled adult wings, with only sporadic sensory bristles (Fig. S2B), indicating disrupted patterning and growth. Overexpression of an aPKC transgene did not induce any visible phenotype (Fig. S2C), and was able to rescue the *aPKC-RNAi* phenotype, confirming the phenotype seen from the RNAi was directly a result of loss of *aPKC* (Fig. S2D). The knockdown of *flfl* via RNAi or over expression of the wild type *flfl* transgene had no affect on the *aPKC-RNAi* phenotype (Fig. S2E, F). Importantly, expression of *N^nucl^* was unable to rescue the crumpled wing from *aPKC* knockdown, indicating aPKC is most likely not a direct upstream regulator of Notch in the wing imaginal disc (Fig. S2G). In addition to this, overexpression of *aPKC* had no effect on the *flfl-RNAi* vein thickening compared to *flfl-RNAi* alone (Fig. 4D, S2H). Although aPKC is involved with PP4 and Notch signaling in ACD (Sousa-Nunes et al., 2009; Zhang et al., 2016), it does not appear to be directly involved in Notch signaling and pattering in the developing wing imaginal disc.

## Discussion

An *in vivo* RNAi screen initially identified three components of the PP4 enzyme complex as modulators of endogenous Wg signaling during wing development (Swarup et al., 2015). The PP4 complex in *Drosophila melanogaster* has been implicated in many signaling pathways, cellular functions and developmental processes, yet its role in regulating Wg signaling was previously uncharacterized. In this study, we revealed that the effect on Wg pathway was at the level of expression of the Wg ligand, by promotion of Notch signaling. Notch signaling regulates the precise expression of Wg in the cells of the D/V boundary of the wing imaginal disc. This expression is required for specification of the wing margin and bristle structures (Couso et al., 1994; de Celis et al., 1996; Neumann and Cohen, 1996).

A partial knockdown of Flfl, PP4-19C and PPP4R2 by RNAi in cells along the A/P boundary of the wing imaginal discs was able to effectively reduce Wg target genes, yet did not induce elevated levels of JNK-mediated cell death as previously reported (Huang and Xue, 2015). Cell death was inducible upon stronger expression of the RNAi using *act-Gal4* with heat-shock inducible flip-out clones (data not shown). The knockdown of Flfl was further found to reduce expression of the Wg ligand as well as other Notch pathway target genes, implicating PP4 in the Notch pathway. Reduction of PP4 proteins in the D/V boundary all resulted in reduced Wg expression, while their knockdown in neighbouring Dl- and Ser-expressing cells had no effect. This result highlights that the PP4 complex acts in boundary cells to regulate Notch-dependent gene expression.

Previously PP4 has been indirectly associated with Notch signaling in Drosophila for its involvement in asymmetrical cell division (ACD) of the developing neuroblasts (Sousa-Nunes et al., 2009; Zhang et al., 2016). While PP4 acts in concert with aPKC and Notch signaling to drive proper ACD, we were unable to identify a role for aPKC in Notch signaling in the epithelial cells of the wing imaginal disc. However, further genetic interaction experiments also identified that PP4’s involvement in Notch signaling in the wing imaginal disc appears to be independent of PP4’s role in cell cycle progression and tissue growth. These results demonstrated that although both PP4 and Notch are required for cell proliferation (Giraldez and Cohen, 2003; Go et al., 1998; Huang et al., 2016), in the wing imaginal disc it is likely not through the same mechanism.

Subsequent genetic interaction studies revealed that most likely Flfl acts to promote Notch through its role in the nucleus. A cytoplasmic form of Flfl could not recue the phenotypes generated by *flfl-RNAi* in the adult wing, while expression of a wildtype transgene could. The required function of Flfl in the nucleus was further bolstered by the fact that the wing phenotype induced by activated nuclear N was partially suppressed by *flfl-RNAi.* As the N^ICD^ enters the nucleus and binds to Suppressor of Hairless and Mastermind to initiate target gene transcription, a multitude of cofactors must be recruited, while others must be removed, from the transcriptional initiation site (reviewed in Bray, 2016). This includes the inhibition of histone deacetylase (HDAC) co-repressor complexes (Kao et al., 1998). PP4 has been previously identified to dephosphorylate and inhibit HDAC activity, while its depletion stimulates HDACs (Zhang et al., 2005). Taken together a possible mechanism for PP4 to promote Notch signaling is through the dephosphorylation of HDACs. This could allow for increased chromatin remodelling, which is needed for the binding of other transcriptional co-factors to ensure full transcriptional initiation of target genes (reviewed Bray, 2016). This is just one possibility, as PP4 may be responsible for the dephosphorylation and modulation of any number of components that cooperate with transcription factors, or regulate the activity of N^ICD^ leading to appropriate target gene expression. Future studies will hopefully address the exact mechanism PP4 plays in promoting nuclear Notch signaling for full expression of target genes like *wg*.

## Methods and Materials

### Fly Strains and Crosses

Fly strains and crosses were raised on standard medium at 25°C unless stated otherwise. *W^1118^* was utilized as wild type. In assays examining genetic interactions between two UAS-driven transgenes, control crosses were performed with *UAS-lacZ* and *UAS-gfp* to eliminate effects caused by the titration of Gal4. The following fly strains were used in this study: *UAS-GFP, UAS-lacZ/TM6B, UAS-dicer, dpp-Gal4, Dll-lacZ, MS1096-Gal4, UAS-aPKC-GFP* (obtained from the Bloomington Drosophila Stock Center), *UAS-GFP, C5-Gal4* (Hugo Bellen), *UAS-flfl Δ 3NLS+2NES (flfl-cyto)* (Gregory Somers), *UAS-flfl* (Zoltan Lipinszki), *wg-lacZ/CyO and UAS-N[nucl]* (Spyros Artavanis-Tsakonas), *UAS-flfl-RNAi* (VDRC 24143,103793), *UAS-PP4-19C-RNAi* (VDRC 25317, 103317, 43250), *UAS-PPP4R2R-RNAi* (VDRC 25445, 105399), *UAS-aPKC-RNAi* (VDRC 2907,105624) [obtained from the Vienna Drosophila Resource Center (Dietzl et al., 2007)], *hh-Gal4/TM6B, UAS-wg-RNAi, wg-Gal4* (ND382) and *en-Gal4,UAS-GFP* (gifts from Konrad Basler, Institute of Molecular Life Sciences University of Zurich, Switzerland).

Crosses involving *C5-Gal4* were performed at 29°C to induce maximal Gal4 expression in the developing wing disc.

### Immunofluorescence of wing imaginal discs

Third-instar larvae were dissected in phosphate-buffered saline (PBS). Wing imaginal discs were fixed in 4% paraformaldehyde at room temperature for 20 minutes and then washed 3 times for 5 minutes in PBS. Discs were blocked [2% BSA diluted in PBS 0.1% Triton X-100 (PBST)] for 45 minutes at room temperature, followed by incubation with primary antibodies (diluted in block) overnight at 4°C. Tissue was then washed 3 times in PBST and incubated with secondary antibodies (diluted in block) at room temperature for 1.5 hours. Tissue was then washed a final three times in PBST and mounted in a 70% glycerol solution. The following primary antibodies and dilutions were used: mouse anti-β-galactosidase (1:2000 Promega), mouse anti-Wg (1:100 DSHB), mouse anti-Cut (1:50 DSHB), mouse anti-N^ICD^ (1:50 DSHB), mouse anti-Delta (1:50 DSHB), mouse anti-Arm (1:50 DSHB), rabbit anti-cleaved Caspase 3 (1:100 Cell Signaling), rabbit anti-PH3 (1:200 Cell Signaling), guinea pig anti-Sens (1:500, a gift from Hugo Bellen, Dept. of Molecular and Human Genetics, Baylor College of Medicine, USA), mouse anti-Dll (1:300, a gift from Ian Duncan, Dept. of Biology, Washington University in St. Louis, USA). Secondary antibodies (Jackson ImmunoResearch) were used at a 1:200 dilution.

### Adult wing mounting

Adult wings were dissected in 95% ethanol followed by mounting in Aquatex (EMD Chemicals Inc.) A minimum of 8 wings were mounted per genotype for analysis.

### Imaging, Analysis and Quantification

Fluorescent images were taken with a Nikon A1R laser scanning confocal microscope and processed using Adobe Photoshop CS6. Adult wings were imaged with an Axioplan 2 microscope. Adult wing and wing disc areas were quantified using ImageJ software. PH3 cell counts were performed using the ImageJ plugin, “Cell Counter”. To compare PH3 positive cell counts per genotype, counts were converted as a ratio of PH3 cells/area, then analysed as experimental condition over control tissue. Significance between groups was assessed by oneway analysis of variance (ANOVA), and *p<0.01* was considered significant unless stated otherwise.

## Acknowledgements

We are grateful to Jessica Blaquiere for technical assistance in the early stage of this study. We also thank Zoltan Lipinszki, Gregory Somers, Spyros Artavanis-Tsakonas, Rita Sousa-Nunes, Hugo Bellen, Juergen Knoblich, Konrad Basler, James Skeath, the Bloomington *Drosophila* Stock Centre, the Vienna *Drosophila* RNAi Centre and the Developmental Studies Hybridoma Bank for providing fly strains and antibodies.

## Competing Interests

The authors declare that they have no competing interests.

## Author contributions

E.T.H., T.P.-S. and E.M.V. designed the experiments, E.T.H., T.P.-S., F.S. and E.M.V. performed the experiments, and E.T.H., T.P.-S. and E.M.V analyzed the data and wrote the manuscript.

## Funding

This work was supported by an operating grant from the Canadian Institutes of Health Research (CIHR) [grant number FRN 133522].

## Hall, Pradhan-Sundd *et al.* - Supplemental Information

**Supplemental Figure 1.**
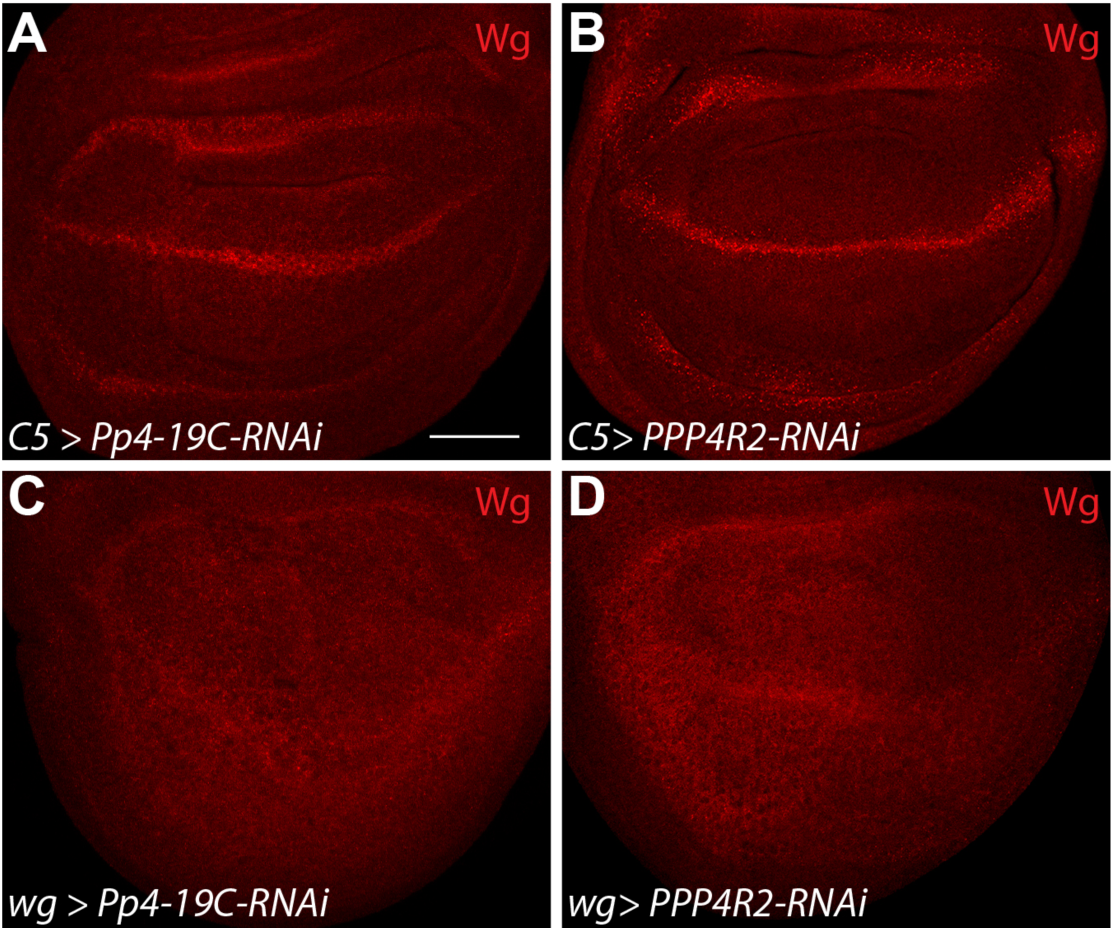
PP4 subunits promote Notch signaling in the Notch receiving cells. (A, B) *C5-Gal4* expressing *Pp4-19C-RNAi* or *PPP4R2-RNAi* along the D/V boundary flanking cells of the wing pouch did not affect Wg protein levels or patterning. (C, D) *wg-Gal4* expression of *Pp4-19C-RNAi* or *PPP4R2-RNAi* resulted in strong reduction of Wg in the wing disc. Scale bar: 50 μm.

**Supplemental Figure 2.**
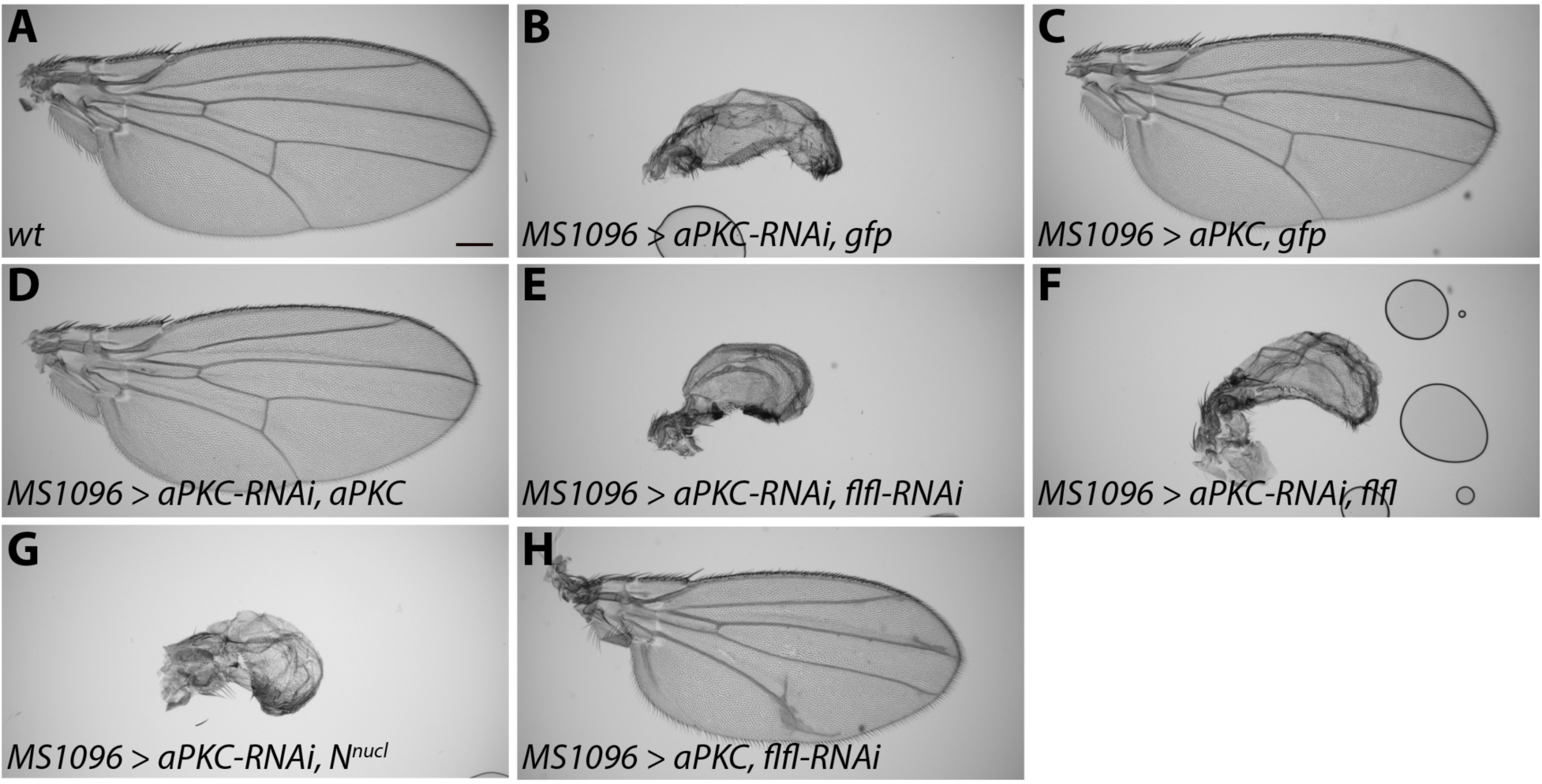
(A) aPKC does not promote Notch signaling in the Drosophila wing. Adult wild-type wing. (B) Knockdown of aPKC with RNAi throughout the entire wing induces small crumpled, blistered, and malformed wings. (C-D) Over expression of an aPKC transgene induces no visible phenotype (C), but can fully rescue the effects of *aPKC-RNAi* (D). (E-G) The knockdown of Flfl (E), or its over expression (F) had no affect on the *aPKC-RNAi* phenotype. N^nucl^ was also unable to rescue the *aPKC-RNAi* phenotype (G). (H) Over expression aPKC does not affect the *flfl-RNAi* phenotype. Scale bar: 300 μm.

